# Drivers of human gut microbial community assembly: Coadaptation, determinism and stochasticity

**DOI:** 10.1101/501940

**Authors:** Kaitlyn Oliphant, Valeria R. Parreira, Kyla Cochrane, Emma Allen-Vercoe

**Author notes:** **Corresponding author** – Kaitlyn Oliphant, University of Guelph, 50 Stone Road East, Guelph, ON, Canada N1G 2W1.

## Abstract

Microbial community assembly is a complex process shaped by multiple factors, including habitat filtering, species assortment and stochasticity. Understanding the relative importance of these drivers would enable scientists to design strategies initiating a desired reassembly for *e.g.*, remediating low diversity ecosystems. Here, we aimed to examine if a human fecal-derived defined microbial community cultured in bioreactors assembled deterministically or stochastically, by completing replicate experiments under two growth medium conditions characteristic of either high fiber or high protein diets. Then, we recreated this defined microbial community by matching different strains of the same species sourced from distinct human donors, in order to elucidate whether coadaptation of strains within a host influenced community dynamics. Each defined microbial ecosystem was evaluated for composition using marker gene sequencing, and for behaviour using ^1^H-NMR based metabonomics. We found that stochasticity had the largest influence on the species structure when substrate concentrations varied, whereas habitat filtering greatly impacted the metabonomic output. Evidence of coadaptation was elucidated from comparisons of the two communities; we found that the artificial community tended to exclude saccharolytic Firmicutes species and was enriched for metabolic intermediates, such as Stickland fermentation products, suggesting overall that polysaccharide utilization by Firmicutes is dependent on cooperation.

## Introduction

A critical knowledge gap in the field of microbial ecology is understanding the relative contribution of the forces that drive microbial community assembly. Uncovering this information would facilitate the development of rationally-designed strategies to remediate microbial communities exhibiting undesirable functionality or successive progression after a perturbation. Such forces have been proposed to include environmental selection, historical contingency, dispersal limitation and stochasticity[1]. Environmental selection additionally encompasses several distinctive sub-factors that yield well to manipulation or measurement for predictions, including niche availability (*i.e.*, habitat filtering) and microbial interactivity (*i.e.*, species assortment and coadaptation)[2–4]. Bioreactors present a promising strategy for studying the importance of such drivers, because culture conditions within them can be tightly controlled, and the use of defined microbial consortia can additionally serve to not only deconvolute the system but also to allow for manipulation to address each factor individually[5, 6]. Bioreactors are also currently utilized for both research and industrial processes[7–13], and thus observing and quantifying the contribution of these ecological forces on community assembly is in itself useful information for these applications.

The human gut microbial ecosystem (*i.e.*, human gut microbiota) is a suitable testing ground for ecological theory. This ecosystem is known to be critical to health and well-being[14–16], with alterations in both community structure and function reported in several GI disorders[17, 18]. Proper succession of the human gut microbiota during infancy and childhood is also essential to proper development and education of the immune system[19], with such deviations again associated with later onset of autoimmune conditions[20–22]. Thus, therapeutics targeting the human gut microbiota have been trialed in such cases, including probiotics and fecal microbiota transplantation. However, the results of such clinical trials have been mixed[23–25], with several factors having been found to influence outcomes, including dosage, number of strains/donor, and diet. Clearly, the availability of an ecological theoretical framework to contextualize the environmental and microbial constitution would improve the design process of these ecosystem interventions. Further, bioreactor-based models such as the Simulator of Human Intestinal Microbial Ecosystem (SHIME) system[26, 27] and single-vessel units[13], are popular methodological approaches for investigating human gut microbial ecology, due to their replicability, sample yields, cost and lack of ethical constraints.

Thus, in our study, we aimed to quantify the relative impact of two of the drivers of microbial community assembly, environmental selection and stochasticity, in terms of both compositional species structure and metabolic behaviour, through use of a defined microbial community derived from a human fecal sample, and single-vessel bioreactor-based models. For the evaluation of environmental selection, we chose to utilize different medium formulations that replicate a high fiber, low protein and a high protein, low fiber diet, as diet has been proposed to be the dominant environmental influencer acting on the human gut microbiota[28]. We additionally scrutinized the two sub-factors of environmental selection, habitat filtering and species assortment, by including a second defined microbial community matching the species constitution of the first, but where each bacterial strain was sourced from a unique human donor. Therefore, the diets would be a representation of habitat filtering, whereas the distinctive communities would model species assortment or coadaptation. To control for co-adaption to the dietary condition that could occur during the initial assembly, we additionally introduced a dietary change after allowing sufficient time for community equilibration to measure the response to a relevant perturbation. Finally, for the evaluation of stochasticity, we assessed the reproducibility of replicates using several multivariate statistical methods to explore steady-state community dynamics.

## Materials and Methods

### Creation of defined microbial communities

Two defined microbial communities were created to examine the effects of coadaptation on the dynamics of community assembly. The first community served as a control, in which all bacterial strains were derived from the same fecal sample. The second community was constructed to match the species composition of the first (as determined by aligning the 16S rRNA genes from each respective pair), but with each bacterial strain sourced from a unique donor’s fecal sample. The isolation methods and donor description of the control community (CC) is described in Petrof *et al.*[29]; however, additional species from this isolation round were added to improve the diversity of the formulation (**Table S1**). The same isolation techniques were utilized to derive the microbial strains for the second, ‘artificial’ community (AC). The donors or international culture collections used to source each species of the AC are indicated in **Table S1**.

Genomic DNA (gDNA) isolated from each strain was individually 16S rRNA gene sequenced using an Illumina MiSeq instrument (Illumina Inc., Hayward, CA, USA) in order to use the high read count output as a method to interrogate the purity of each sample. The gDNA was first extracted from each strain following the protocol described in Strauss *et al.*[30], and the library preparation, sequencing and data processing were then conducted as described in the *16S rRNA based compositional profiling* section below. An average read depth in the 10^3^ range was typically achieved when sequencing single strains. The strains were decidedly pure when all amplicon sequence variants (ASVs) that could not be attributed to the target species were of low abundance (< 1%) and could be accounted for as sample cross-contamination through referencing sample blanks. Any strains that were not pure were subjected to serial dilution to extinction, as a method to improve isolation, and were then re-evaluated by the above technique.

### Bioreactor operation

A 500 mL Multifors bioreactor system (Infors AG, Bottmingen/Basel, Switzerland) was inoculated with the defined microbial communities and operated as a model of the human distal colon as previously described[31]. The feed medium was designed to replicate two dietary conditions, high fiber, low protein (HF) and high protein, low fiber (HP), based upon the formulation in Marzorati *et al.*[32] but modified to accommodate a single-vessel system as in McDonald *et al.*[13] (**Table S2**). The experimental design is depicted in **Figure 1**.

**Figure 1.**
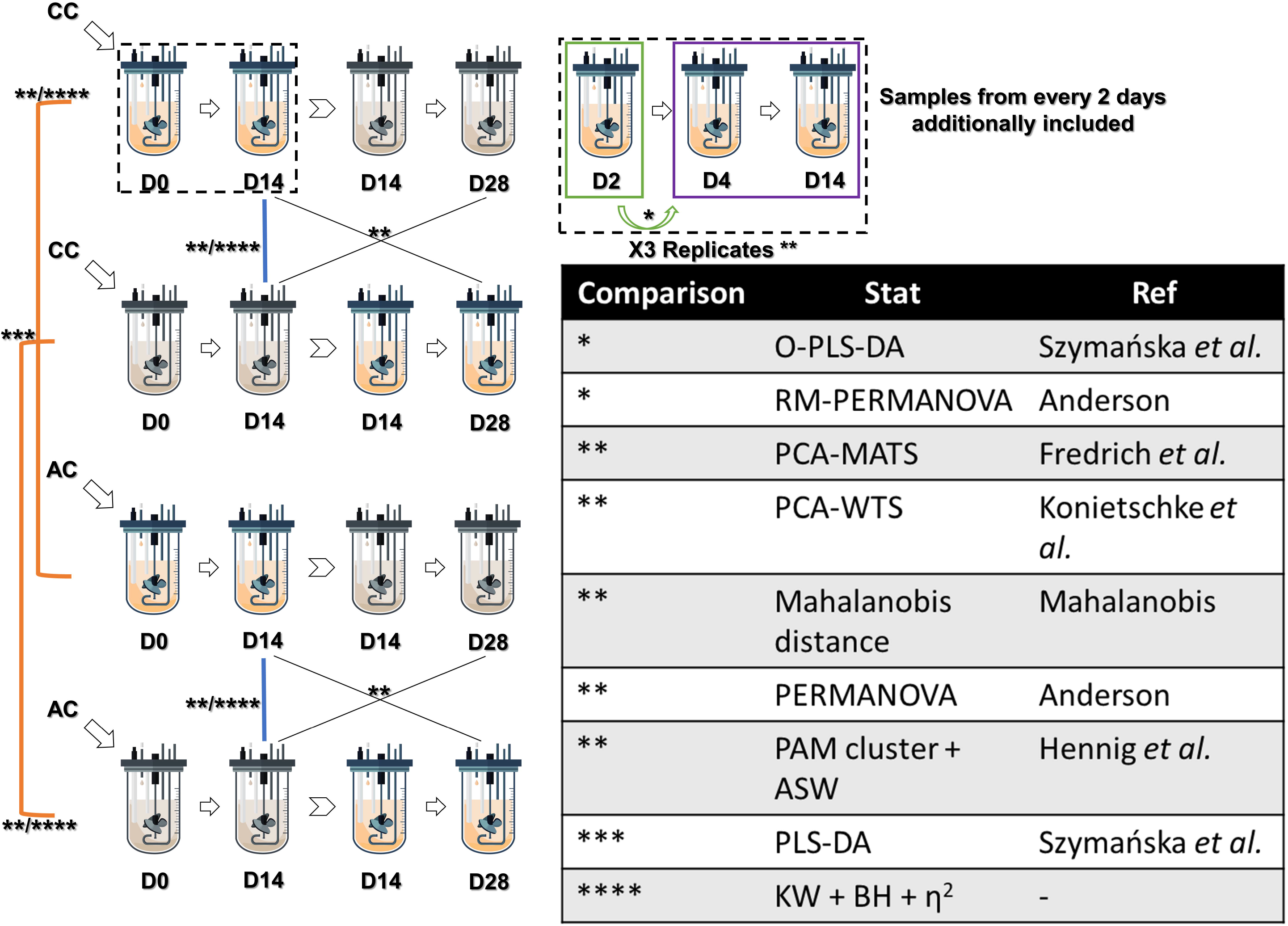
Experimental design of bioreactor runs and statistical testing. Bioreactors were either inoculated with the control microbial community (CC) or the artificial microbial community (AC). Bioreactors were either fed a high fiber, low protein medium or a high protein, low fiber medium, which are distinguished by colour in the figure. After allowing a 14-day equilibration, the medium formulations were changed, and then the bioreactors were run for an additional 14 days. Sampling occurred every 2 days, in which 2 × 2 mL were taken and stored at -80 °C for 16S rRNA sequencing and ^1^H-NMR based metabonomics, respectively. The box with the dotted line showcases this sampling schematic. Both the composition, *i.e.*, species structure, and metabolic signature, *i.e.*, binning data, were utilized for all comparisons. Comparison 1 (*) was done to resolve steady-state; samples were first divided into two groups, ‘early’ and ‘late’, and moving window analysis was done using a valid orthogonal partial-least squared discriminant analysis (O-PLS-DA) model, as determined by permutation testing described in Szymańska *et al.*[80] using the R software (version 3.5) package ropls 1.12. As well, the p-value computed from a PERMANOVA of the Euclidean distance matrix as described in Anderson[81] using the R package vegan version 2.5.2, with permutations restricted in a repeated measures design [81, 82]. Only steady-state time points were used for subsequent comparisons. The remaining comparisons (**) conducted were within replicates of the same condition, between media (controlling for microbial community), between microbial communities (controlling for medium), and between the treatment versus starting diet. These tests included dimension reduction by principal component analysis (PCA) then the Wald-type statistic (WTS) for multivariate data as according to Konietschke *et al.*[83] and the modified ANOVA-type statistic (MATS) as according to Fredrich and Pauly[84]. Resampling was achieved with a parametric bootstrap approach via the R packages ropls and MANOVA.RM version 0.3.1. Additionally, the pairwise Mahalanobis distances[85, 86] were calculated using R package HDMD version 1.2. PERMANOVA of the Euclidean distance matrix was also computed as described in Anderson[81] using the R package vegan. Additionally, PAM clustering was conducted, selecting the best solution from the average silhouette widths (ASWs)[87] using the package fpc version 2.1.11. Metabolites of interest were selected from partial least squared discriminant analysis (PLS-DA, ***) of all samples grouped by the microbial community and medium combination via variable importance of projection (VIP) scores (> 1) using R package ropls. Statistically significant differences between individual taxa and metabolites (****) were then determined between media and between microbial communities grown within each medium by a series of Kruskal-Wallis (KW) tests with Benjamini-Hochberg (BH) correction and Dunn’s post-hoc analysis via the package dunn.test version 1.3.5, in addition to the calculation of effect size (η^2^). Only features that remained significant after post-hoc analysis (q-value < 0.05) and had an effect size above 50% were considered significant.

### 16S rRNA based compositional profiling

The gDNA from bioreactor samples was extracted through use of the QIAamp Fast DNA Stool Mini Kit (Qiagen Inc., Germantown, MD, USA) according to the manufacturer’s directions, but with extra steps included to improve cell lysis. Prior to proceeding with their recommended protocol, cells were first pelleted through centrifugation at 14 000 rpm for 15 min at 4 °C. After resuspension in the lysis buffer, 0.2 g of zirconia beads (Biospec Products Inc., Bartlesville, OK, USA) were added, then the samples were bead-beat with a Digital Disruptor Genie (Scientific Industries Inc., New York City, NY, USA) at 3000 rpm for 4 min. The samples were subsequently incubated at 90 °C for 15 min, and finally, ultrasonicated at 120 V for 5 min (Branson Ultrasonics, Danbury, CT, USA). Libraries for sequencing were constructed by a one-step PCR amplification with 400 ng of Nextera XT Index v2 sequences (Illumina Inc.) plus standard 16SrRNA v4 region primers[33] and 2 μL of gDNA template in Invitrogen Platinum PCR SuperMix High Fidelity (Life Technologies, Burlington, ON, Canada). Cycler conditions included an initial melting step of 94 °C for 2 min, followed by 50 cycles of 94 °C for 30 s, annealing temperature for 30 s and 68 °C for 30 s, with a final extension step of 68 °C for 5 min. The annealing temperature comprised of a 0.5 °C increment touch-down starting at 65 °C for 30 cycles, followed by 20 cycles at 55 °C. The PCR products were subsequently purified using the Invitrogen PureLink PCR Purification Kit (Life Technologies) according to the manufacturer’s directions. Normalization and Illumina MiSeq sequencing was carried out at the Advanced Analysis Center located in the University of Guelph, ON, Canada.

The obtained sequencing data was processed using R software version 3.5 with the package DADA2 version 1.8, following their recommended standard protocol[34]. Classification to the genus level was additionally carried out via DADA2 using the SILVA database[35] version 132. Classification to the species level, however, was conducted by uploading the ASVs to NCBI BLAST (https://blast.ncbi.nlm.nih.gov) and selecting the identification with the highest percentage and lowest e-value, while cross-referencing with the known species constitution of the defined microbial communities. The data was then denoised by adding the ASVs that returned identical species classifications together, and after which the ASVs that equated to < 0.01% total abundance across all samples were removed. Finally, the data was normalized by center-log ratio transformation through use of the package ALDeX2 version 1.12, taking the median Monte-Carlo instances as the value[36]. The statistical analysis of this data is described in **Figure 1**.

### ^1^H-NMR based metabonomics

Sample preparation, ^1^H-NMR spectral acquisition and processing, and profiling of metabolites was conducted as previously described[31]. A Bruker Avance III 600 MHz spectrometer with a 5 mm TCI 600 cryoprobe (Bruker, Billerica, MA, USA) at the Advanced Analysis Center located in the University of Guelph, ON, Canada was utilized for spectral acquisition. Spectra were collected at a sample temperature of 298 K. The data was analyzed using both an untargeted spectral binning and targeted metabolite profiling approach with the Chenomx NMR suite 8.3 (Chenomx Inc., Edmonton, AB, Canada). For spectral binning, the default parameters of 0.04 ppm sized bins along the 0.04 – 10 ppm region of the spectrum line with omission of water (4.44 – 5.50 ppm) and normalization by standardized area (fraction of the chemical shape indicator, DSS) were implemented. For metabolite profiling, target regions of interest in the spectra were selected by partial least squared discriminant analysis of the spectral binning data (**Figure 1**). Metabolite identifications were then based on the best fit for the peak regions with the available libraries of compounds. The libraries included both the internal set included with the Chenomx software suite, and the downloaded HMDB[37] set release 2. The statistical analysis of both data sets is described in **Figure 1**.

## Results

### Determination of microbial community ‘steady-state’ stability and replicate reproducibility

For this work, it was essential to first determine at which day the bioreactor-grown microbial communities had achieved a stable equilibrium, referred to as ‘steady-state’, in order to make subsequent comparisons. The CC reached steady-state both compositionally and metabolically by day 4 (**Table S3**). Upon dietary change from HF to HP, the microbial community was compositionally similar from the first time point (day 16) until the end of the run (day 28), however, metabolic stability was not reached until day 18. This latter observation is in line with the calculated amount of time it would take for the bioreactor to shed the excess 2000 mg/mL concentration of fiber from lingering fiber-rich medium following medium change to HP composition, which is 4 days post medium change (**Figure S1**).

Results obtained running the AC under the same conditions were much more variable (**Table S3**). Steady-state was reached for this community both compositionally and metabolically by day 2 in the HF medium, with no significant differences observed at the first time point of day 16 after the change in medium formulation. In the HP medium, however, the AC followed the patterns of the CC more closely than in the HF medium, except for taking longer to reach initial metabolic stability (6 days compared to 4 days for the CC).

Next, the reproducibility between replicates of the same condition was evaluated for both the normalized sequence count and ^1^H-NMR spectral binning data. Statistically significant differences between replicates were found within all conditions for both data sets (**Table S4**). The pairwise Mahalanobis distances ranged in magnitude from 4.3 to 10.6 for the compositional data and 9.5 to 19.1 for the metabolic data by PCA. The untargeted PAM clustering approach failed to recapitulate the expected patterns, *i.e.*, clustering the samples by replicate. The ASWs ranged in magnitude from 0.23 to 0.28 for the compositional data and 0.13 to 0.24 for the metabolic data. Stochastic variation was thus clearly present across each replicate experiment, and thus in order for the changes induced by an introduced environmental pressure to be deemed statistically significant overall, we defined this to mean that it must exceed the within-condition replicate separation (**Figure 2**). This could be evaluated by overlap of ellipses and magnitude of the Mahalanobis distances for the PCA approach, and recapitulation of the expected clusters by untargeted PAM clustering with improved ASWs for the Euclidean distance approach.

**Figure 2.**
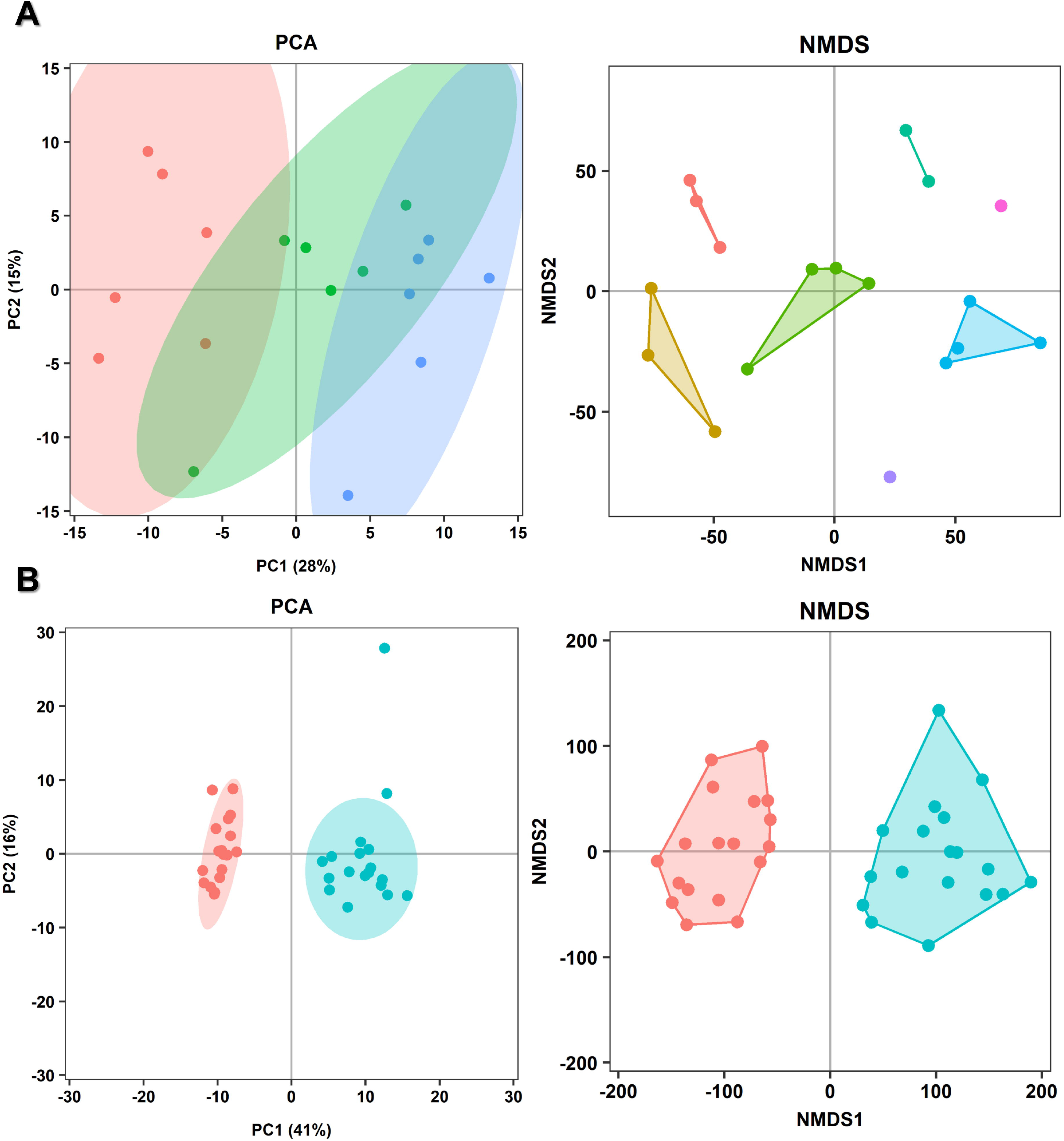
Analysis of overall statistically significant differences in the ^1^HNMR spectral binning data obtained for the control community. Panel **A** depicts samples from days 2 – 14 of the three bioreactors fed the high fiber medium formulation, grouped by replicate (1 – 3). Panel **B** depicts samples from days 2 – 14 of all six bioreactors grouped by medium formulation (high fiber and high protein). The **left** panel is the result of principal component analysis (PCA) conducted in R software version 3.5 by the package ropls version 1.12, with plots, including ellipses assuming the multivariate t distribution, drawn by the package ggplot2 version 2.2.1. The **right** panel is the result of non-multidimensional scaling (NMDS) of Euclidean distance matrices conducted by the package vegan version 2.5.2, with plots, including the best solution of untargeted partitioning around medoids clustering determined by average silhouette width using the package fpc version 2.1.11, drawn by the package ggplot2. Panel **B** presents a superior separation than panel **A**, indicating that the effect of diet exceeds stochasticity.

### Microbial community response to dietary changes

With the decided criteria from the above objective, *i.e.*, the between-group Mahalaonbis distance exceeding the found within-group Mahalaonbis distance and untargeted PAM clustering recapitulating the expected pattern of clustering samples by diet, neither community altered its composition in response to dietary change (**Table S4**; **Figure 2**). However, the dietary change did elicit a clear significant difference in the metabolite profile of both communities (**Table S4**; **Figure 2**). The untargeted PAM clustering approach reliably recapitulated two clusters, each containing only the samples of one specific diet, as the best solution. The ASWs also exceeded the values obtained from the clustering solutions within conditions (replicates), at 0.29 and 0.25 for the CC and AC respectively. The Mahalanobis distance derived from PCA was also larger than the maximum value obtained between replicates of the CC, *i.e.*, 19.3 compared to 18.1. For the AC, however, the result was less clear, as the Mahalanobis distance was less than the maximum value obtained between replicates, *i.e.*, 17.6 compared to 19.1. Finally, there was no significant difference between communities that were originally cultured in one medium compared to those eventually cultured in the same medium following a period of culture in a different medium suggesting that within-experiment adaption was not a confounding factor (**Table S4**).

Based upon the above results, we determined which features (individual taxa and metabolites) were statistically significantly different between the medium formulations of which the communities were initially grown. The normalized sequencing data was used to evaluate shifts in taxonomic abundance (**Table S5**), and as expected, no statistically significant differences were found (data not shown). The ^1^HNMR metabolite profiles (**Table S6**), however, revealed > 15 metabolites that exhibited significant changes in concentration in both the CC and AC (**Table S7**). The concentrations of short-chain fatty acids (SCFAs) and select metabolites of interest that were significantly different between growth medium conditions are depicted in **Figure 3** and **Figure 4** respectively. Most of these alterations were identical between each community, including increased concentrations of several amino acids and their specific fermentation by-products, a lower concentration of methanol and a higher concentration of uracil in the HP medium. Community-specific deviations included increases in concentration of several amino acids and succinate, and a decrease in concentration of valerate for the CC in the HP medium, whereas the AC had a higher concentration of isobutyrate and a lower concentration of glyoxylic acid.

**Figure 3.**
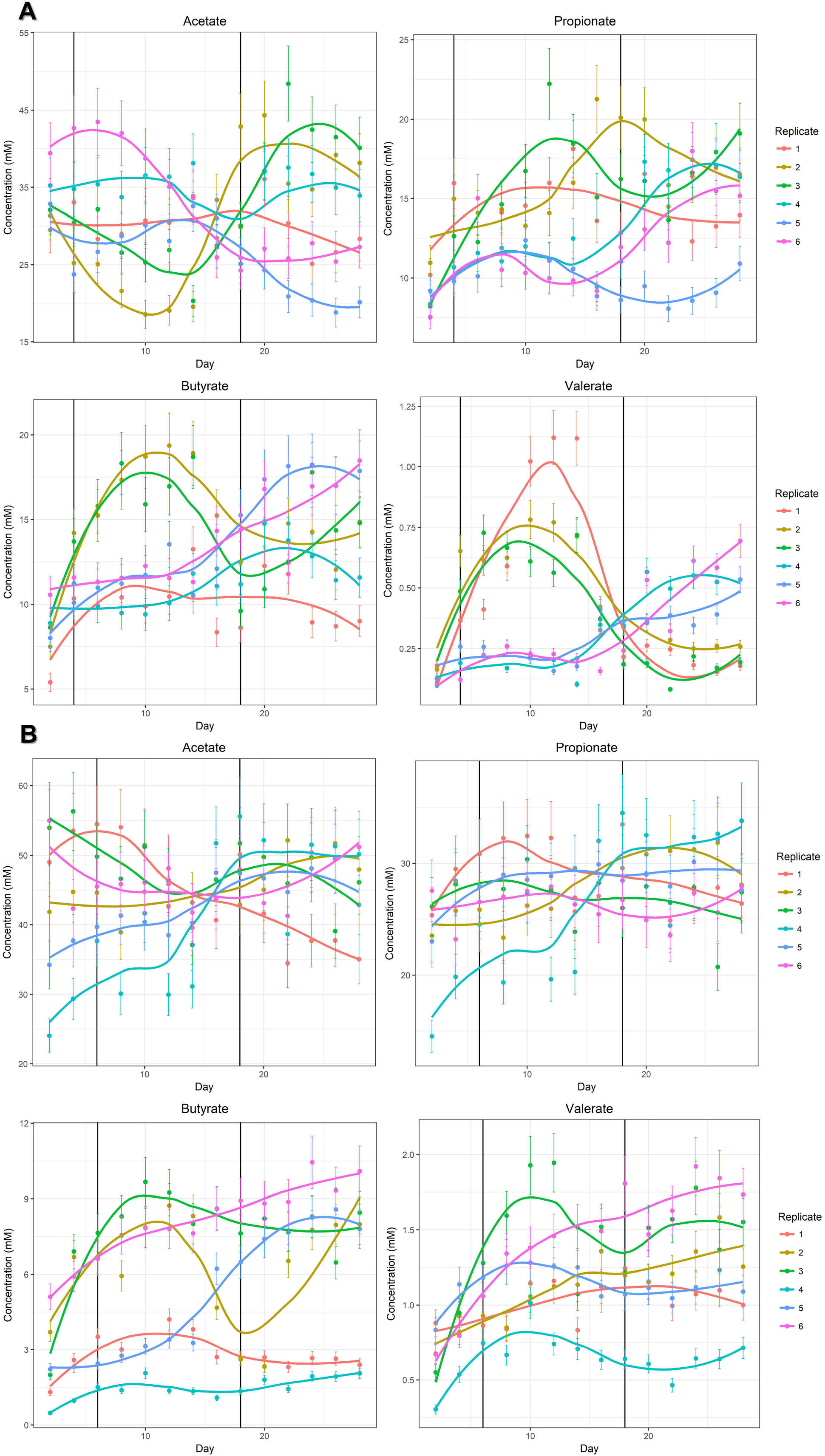
Concentrations of short-chain fatty acids determined by ^1^HNMR metabolite profiling in bioreactor samples over time. Replicates 1 – 3 are the communities that were initially grown in the high fiber medium formulation, whereas replicates 4 – 6 are the communities that were initially grown in the high protein medium formulation. Panel **A** depicts the control community, and panel **B** depicts the artificial community. Plots were drawn in R software version 3.5 by the package ggplot2 version 2.2.1, with 10% error bars representing the expected amount of technical measurement inaccuracy[88].

**Figure 4.**
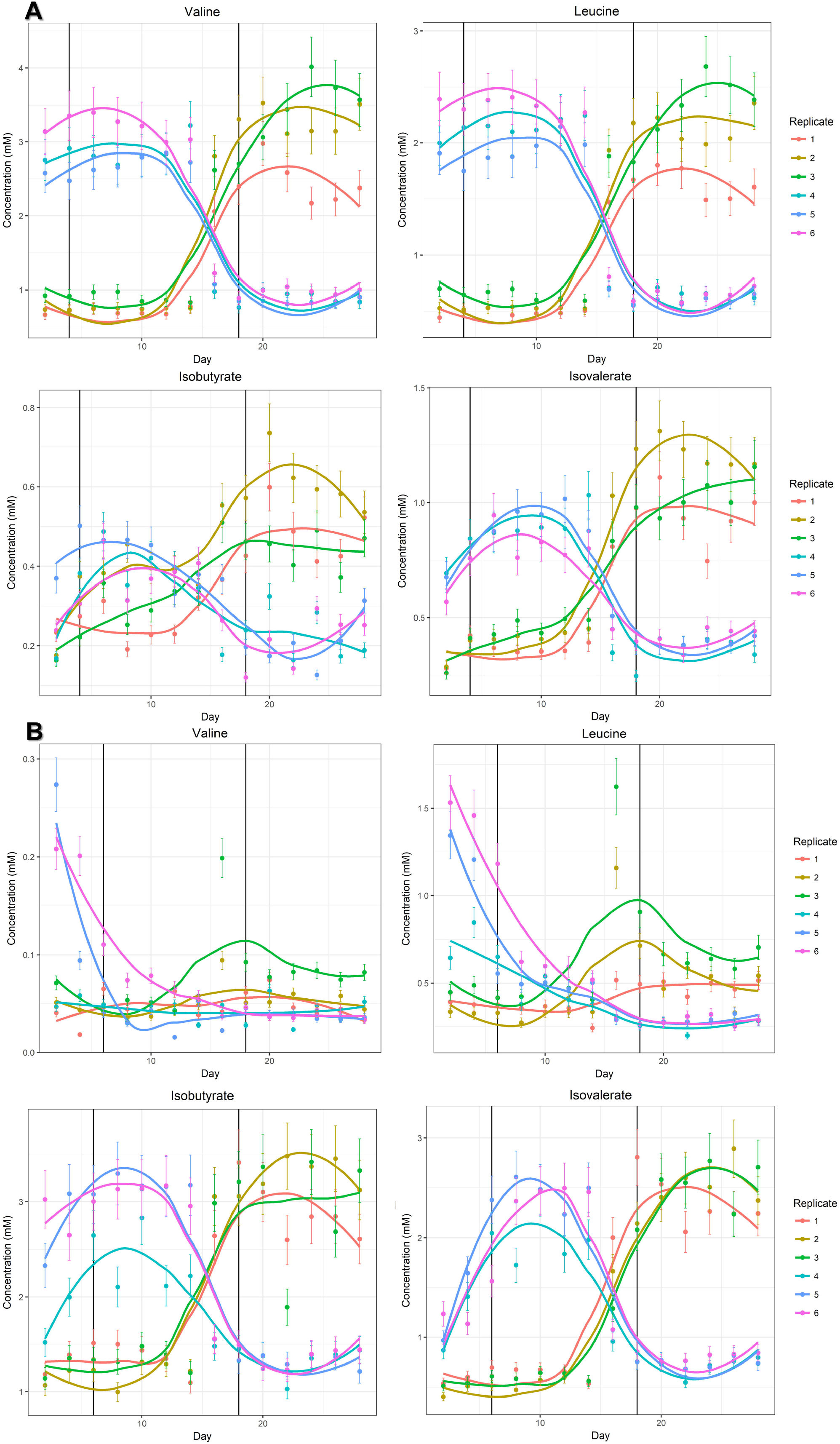
Concentrations of select metabolites that were statistically significantly different between medium formulations or communities as determined by ^1^H-NMR metabolite profiling in bioreactor samples over time. Replicates 1 – 3 are the communities that were initially grown in the high fiber medium formulation, whereas replicates 4 – 6 are the communities that were initially grown in the high protein medium formulation. Panel **A** depicts the control community, and panel **B** depicts the artificial community. Plots were drawn in R software version 3.5 by the package ggplot2 version 2.2.1, with 10% error bars representing the expected amount of technical measurement inaccuracy[88].

### Effect of coadaptation on microbial community structure and behaviour

Finally, the differences between the CC and AC in both media were evaluated overall and between individual taxa and metabolites as above, to determine if potential coadaptation impacted community composition or behaviour. With the set criteria, there were no significant differences between the overall compositional nor metabolite landscape (**Table S4**). However, there were several significant differences between individual taxa and metabolites. For the taxa, *[Eubacterium] rectale* (HF/HP; p-value = 5E-6/4E-4; effect size = 74%/62%)*, Faecalicatena fissicatena* (HF/HP; p-value = 2E-5/6E-4; effect size = 60%/54%) and *Coprococcus comes* (HF only; p-value = 1E-5; effect size = 65%) were significantly altered. Upon examining the raw sequence counts (**Table S5**), it was observed that both *E. rectale* and *C. comes* were virtually undetected in the AC, with the latter likely only reaching statistical significance in the HF condition due to its relatively higher abundance in the CC. On the other hand, *F. fissicatena* was present in both communities, but at a lower abundance in the CC. For the metabolites, several amino acids, organic acids and uracil had increased concentrations in the CC, whereas branched-chain fatty acids reached higher concentrations in the AC (**Table S6**; **Table S7**; **Figure 4**).

## Discussion

Understanding the drivers of community succession is not only useful for researchers utilizing technologies to simulate microbial ecosystems, such as bioreactors, but also can translate to real world applications. For the human gut microbiota, such an understanding could assist in the design of therapeutic strategies aiming to ameliorate abnormalities exhibited in GI disorders, or the building of predictive tools to project changes over time (for example, during infant development). Here, we examined the relative impact of environmental selection and stochasticity on gut microbial community succession through use of HF and HP medium formulations. Further, we differentiated the effects of habitat and member coadaptation within environmental selection, by creating both a microbial community derived from a single fecal sample and an equivalent community (same or highly similar species) with each member sourced from individual donor fecal samples.

First, it was essential to determine the point at which the microbial communities form a stable equilibrium, known as ‘steady-state’. Removing time level variation is important when using bioreactor-based models, as it eliminates technical artifact bias that can confound results. We found that for the CC, steady-state was reached by day 4. This result is much earlier than has been suggested by other studies using SHIME[26, 27] or single-vessel systems[13]. There are several potential reasons for this discrepancy. First, this previous work has used observational techniques, such as moving window analysis or Unifrac clustering, to determine the day at which the microbial community had stabilized. These methods do not statistically validate the variation, resulting in overestimation of the value. Second, steady-state tends to be measured within each replicate, instead of assessed as a dataset. Microbial community dynamics between replicates were not found to be 100% reproducible. Therefore, ‘steady-state’ can ultimately be defined as the point at which time level variation no longer exceeds the extent of replicability, as upon reaching this range statistical tests would no longer be confounded by this technical property. Finally, biological differences could also attribute to this low value, since our experimental communities were of a reduced complexity compared to a typical fecal ecosystem (for example, since there are fewer interaction types and processes in our defined experimental communities compared to fecal communities, equilibrium is achieved more quickly when projected by relevant mathematical models, such as Lotka-Volterra[38]). Of note, we recommend our approach of utilizing whole sequence count and spectral binning data, as opposed to individual taxa and metabolites, to gauge steady-state for a complex microbial community. Not all metabolites stabilized in concentration at the same time, and the rate at which individual metabolites reached stable concentrations was not identical between replicates. Particularly, the metabolites most frequently measured in human fecal-associated communities due to their importance and dominant concentrations, the SCFAs[39, 40], often stabilized faster than other metabolites *e.g.*, amino acid-derived fermentation by-products (**Figure 3**; **Figure 4**).

The property of significant variation between replicates of the same microbial community under identical conditions is not a new observation for bioreactors[7–11]. It has been termed ‘multistability’, which occurs when numerous entities have non-linear interactions or feedback loops, and thus multiple alternative stable states are able to exist[41]. This phenomenon has been observed both in environmental ecosystems *in situ*[42, 43], including the human gut microbiota, with the best example of it being reported after antibiotic treatment[41, 44, 45]. Therefore, multistability is not an artifact of the modeling technology but rather an accurate representation of this naturally inherent property. Switching between such states can occur either under a gradual external influence or after experiencing a substantial perturbation[41]. Bioreactors thus present a useful tool for quantifying the extent of this stochasticity and determining which factors induce microbial community reassembly through examining within run variability over time. However, it is also important for researchers to ensure that the applied perturbation in their experiment exceeds this inherent variability in order to demonstrate an overall change in microbial community composition or behaviour; simply obtaining a significant p-value is not enough, as such a result can be attained when comparing replicates of identical conditions. We therefore recommend calculating the differences between and within conditions in the Mahalanobis distance of groups after PCA and ASW plus cluster membership after PAM clustering of Euclidean distance matrices (**Figure 2**).

We found that growth medium did significantly alter the microbial community metabolic behaviour but not the composition. The latter result is in line with previous observations that found only minor alterations in microbial community species structure after a dietary change[46–48], and the composition of the human gut microbiota is also reportedly robust in adulthood, with a 60% microbial strain retention rate in a five-year window[49]. Additionally, the metabolic changes we saw were as expected, both proteolysis (higher amino acid concentrations) and amino acid fermentation (higher concentrations of by-products specific to these metabolisms[15, 50]) increased in the HP medium (**Figure 4**). Our resultant lack of difference between the assembled and switched microbial communities growing in the same medium is supported by Wu *et al.*[48], as the authors found that dietary changes only influenced the species structure of the human gut microbiota when maintained for a much longer term as opposed to ten days. We would thus conclude that for the typical duration of bioreactor experiments, it is not necessary to consider microbial community adaptation as a confounding factor.

The control and artificial communities were similar, but not identical to each other, as indicated by the non-significant differences in overall composition and metabolic behaviour. In terms of species structure, both *E. rectale* and *C. comes* failed to integrate into the AC. Both species are saccharolytic and capable of degrading fructans *e.g.*, inulins; *E. rectale* can also utilize starch and xylan-derived polymers[51–53]. Intriguingly, Lozupone *et al.* built a co-occurrence network from fecal metagenomic data collected from 124 unrelated adults and found that *C. comes* co-occurred with *E. rectale*[54]. *E. rectale* in particular is known to require cooperation with other species to utilize resistant starches, as it is incapable of conducting this activity on its own[55, 56]. Lozupone *et al.* found that *Bacteroides* spp. additionally co-occurred with *E. rectale* and *C. comes*, thus the *Bacteroides ovatus* strains in our communities represent potential collaborators[54]. An elegant study conducted by Rakoff-Nahoum *et al.* discovered that a strain of *B. ovatus* produced both membrane-bound and secreted forms of a glycoside hydrolase capable of degrading inulin[57]. When the secreted form of the enzyme was knocked-out, the fitness of *B. ovatus* was not impacted when grown in monoculture but was significantly diminished when grown in a community setting. Tuncil *et al.* additionally demonstrated that different *Bacteroides* spp., namely *Bacteroides thetaiotaomicron* and *B. ovatus*, had reciprocal glycan substrate preferences that were maintained from monoculture to coculture[58]. The combination of these two studies would indicate potential mechanisms of coadaptation within a gut microbial ecosystem, such that species would adapt to occupy unique niches or collaborate to exploit the same niche, at least in terms of polysaccharide consumption. Therefore, it is possible that the *B. ovatus* strain in the AC lacked secretory catabolic enzymes that would have assisted *E. rectale* and *C. comes* in utilizing *e.g.*, fructans, or that the glycan substrate preference of the *B. ovatus* had shifted to consuming the fructans that *E. rectale* and *C. comes* could have otherwise degraded themselves, thus effectively outcompeting them. Alternative explanations for our findings, beyond metabolism, might include, for example, a communication mismatch between strains through via quorum sensing[59] or incompatible antimicrobial defense mechanisms[60]. This result thus not only revealed the presence of a possible, cooperative microbial guild within the CC, but also demonstrated that strain level variation is an important property to consider in studies aiming to examine microbial community cooperation.

In terms of metabolic difference between the studied communities, an intriguing observation was the elevated amount of amino acid fermentation in the AC compared to the CC, as supported by decreased amino acid concentrations and increased concentrations of their specific fermentation by-products[15, 50]. Further, this heightened activity would explain the lower amount of separation between clusters by medium formulation when evaluating overall statistical significance in the AC comparted to the CC. Particularly, there was evidence of Stickland fermentation occurring (higher concentrations of isobutyrate, isovalerate and valerate), a metabolism specific to the Firmicutes (usually the Clostridia) (**Figure 4**)[61, 62]. An interesting finding by Shoaie *et al.* was that when *E. rectale* was cocultured with *B. thetaiotaomicron*, it switched its gene expression from fermenting saccharides to amino acids[63]. The *E. rectale* strain in this case thus likely responded to the introduced competition by occupying a different niche. Together, these results would suggest that polysaccharide utilization by the *Firmicutes* is dependent on a collaborative effort due to this phylum’s more highly specialized metabolisms[64, 65], and in the absence of cooperation, these species will become putrefactive instead. Many GI disorders are characterized by an inherent low diversity[66], and a loss of bacteria from clostridial cluster XIVa, which includes *E. rectale and C. comes*[67, 68]. Further, there is often a concurrent lack of butyrate, produced mainly from fermentation by certain *Firmicutes* members[69], and increased inflammation, which can be promoted by the products of protein fermentation[15, 70, 71]. Therefore, we suggest that an erosion of cooperative interactivity has occurred in these situations, resulting in extinction of particularly dependent species and altered behaviours from those that remain, which exacerbate symptoms.

One limitation to our study was the methods utilized to determine microbial strain purity. Despite our best efforts to confirm purity, we found these techniques to be inadequate at detecting and removing *all* contaminants. Specifically, in the CC, we found that *Akkermansia muciniphila* bloomed in the bioreactors, which was initially unexpected. Upon completing an *Ak. muciniphila* specific PCR[72] on the gDNA collected from each strain, we found it was present in the *Acidaminococcus intestini* stock. This result was unexpected, since *Ak. muciniphila* was completely undetected by marker gene sequencing, and since an average of >10 000 total reads were attained using this method, its presence was indicated at a rate of less than one in every 10 000 cells (**Table S8**). When we completed a re-extraction and re-sequencing of gDNA obtained from *Ac. intestini* cultured on FAA supplemented with mucin, the preferred growth substrate of *Ak. muciniphila*[73], we were then able to detect it; at this point it achieved 10% of the total growth (**Table S8**). To compensate for this unexpected *Ak. muciniphila* load, we added a strain of *Ak. muciniphila* to the AC prior to conducting the bioreactor experiments for this community and then scrutinized the sequencing count data for any other outliers once completed (**Table S6**). We found an unexpected number of reads classified as *Phascolarctobacterium*, and when *Phacolarctobacterium* specific PCRs[74, 75] were conducted on the gDNA of the bioreactor samples and strains, we found *Phascolarcterobacterium faecium* to be present in all six replicates and the *E. rectale* stock of the AC, but not in the CC. Again, it was present at a rate of 1 per 10 000 cells of *E. rectale. P. faecium* did not reach high numbers, however, as it was not statistically significantly increased in the AC when compared to the CC. The fact that it had amounted to above 1% total percentage contribution may be attributed to sequencing error or cross-contamination (**Table S5**). We examined the possibilities as to how its inclusion could confound our experiment. Interestingly, *E. rectale* did not integrate into the AC, so any interactions resulting from coadaptation between it and *P. faecium* were absent. *Ac. intestini* was also able to colonize the CC with a contaminant present at an equivalent amount as *E. rectale*, so we doubt its inclusion would have negatively impacted the ability of *E. rectale* to incorporate into the AC either. *P. faecium* is asaccharolytic and incapable of Stickland fermentation, and instead consumes succinate as a substrate to produce propionate[74, 76]. Therefore, the inclusion of *P. faecium* would not have altered any of our discussion points regarding polysaccharide utilization networks or increased putrefactive activity. The only influence it thus potentially had was the significant decrease in the concentration of succinate and increase in the concentration of propionate in the AC (**Figure 3**). We thus determined that our experiment remained of sufficient validity to answer our hypothesis, and decidedly continued due to a lack of available and validated alternatives for detecting contaminants. Hopefully, new technology that can improve sequencing accuracy, reduce cross-contamination and enhance the obtained number of reads will address this issue in the future; for a review of current next-generation sequencing developments, please refer to Goodwin *et al.*[77]. However, these drawbacks should presently be carefully considered by other researchers.

We have concluded that stochasticity is a property inherent to human gut microbial ecosystems but is exceeded by forces of environmental selection, at least in terms of driving microbial community behaviour. Substrate availability also seems to dictate functionality over cooperative interactivity, but that does not preclude the existence of coadaptation. Our work fits into the observations of previous studies, that indicate microbial community assembly in the human gut is a deterministic process[78], habitat filtering predominates species assortment[2], competitive interactions are more numerous than cooperative ones[79], but microbial guilds covering metabolic modules exist[3, 4]. We have also suggested a methodology to elucidate steady-state, that additional testing is required to determine overall statistically significant differences of a treatment, and that microbial community alterations exhibited in GI disorders could result from a break-down of cooperation. Future work studying microbial interactions should consider strain level variation, and could, for example, compare interactions between strains derived from ‘healthy’ versus low diversity ecosystems.

## Acknowledgements

We would like to acknowledge the Natural Sciences and Engineering Research Council of Canada and Ontario Ministry of Training, Colleges and Universities scholarships to KO for providing funding. Funding was gratefully received from an NSERC Discovery Grant and a National Institutes of Health R33 to EA-V. We are most thankful to our collaborators, Dr. Mike Surette, McMaster University, Canada and Dr. Sydney Finegold, VA Wadsworth Laboratory, USA for the gift of suitable matching strains for our AC community, as well as members of the EA-V lab for providing other needed isolates – Dr. Mike Toh, Michelle Daigneault, Erin Bolte and Dr. Rafael Peixoto.

