## Supplementary figures and images for "Drivers of human gut microbial community assembly: Coadaptation, determinism and stochasticity"

### Figure S1

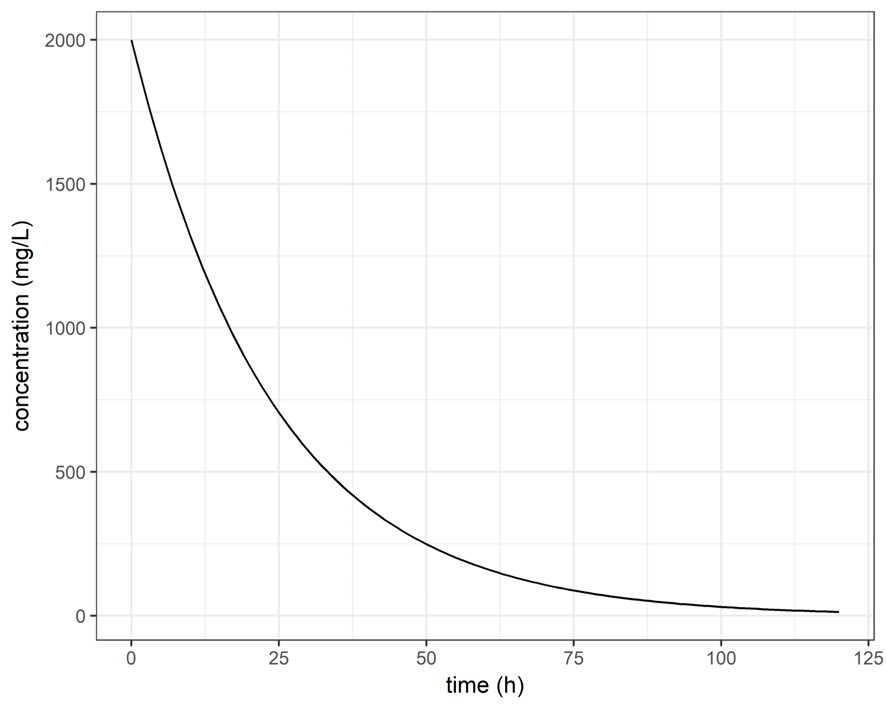
